# Dynamics of morphogen source formation in a growing tissue

**DOI:** 10.1101/2024.03.01.582751

**Authors:** Richard D. J. G. Ho, Kasumi Kishi, Maciej Majka, Anna Kicheva, Marcin Zagorski

## Abstract

A tight regulation of morphogen production is key for morphogen gradient formation and thereby for reproducible and organised organ development. Although many genetic interactions involved in the establishment of morphogen production domains are known, the biophysical mechanisms of morphogen source formation are poorly understood. Here we addressed this by focusing on the morphogen Shh in the vertebrate neural tube. Shh is produced by the adjacently located notochord and by the floor plate of the neural tube. Using a data-constrained computational screen, we identified different possible mechanisms by which floor plate formation can occur, only one of which is consistent with experimental data. In this mechanism, the floor plate is established rapidly in response to Shh from the notochord and the dynamics of regulatory interactions within the neural tube. In this process, uniform activators and Shh-dependent repressors are key for establishing the floor plate size. Subsequently, the floor plate becomes insensitive to Shh and increases in size due to tissue growth, leading to scaling of the floor plate with neural tube size. In turn, this results in scaling of the Shh amplitude with tissue growth. Thus, this mechanism ensures a separation of time scales in floor plate formation, so that the floor plate domain becomes growth dependent after an initial rapid establishment phase. Our study raises the possibility that the time scale separation between specification and growth might be common strategy for scaling the morphogen gradient amplitude in growing organs. The model that we developed provides a new opportunity for quantitative studies of morphogen source formation in growing tissues.

## Introduction

Morphogen gradients are key for patterning of developing tissues. In the last few decades, much work has established the principles by which morphogen gradients form (*1*). In many systems, morphogens form exponential gradients within their target tissues, as a result of morphogen production from a restricted source, non-directional transport and degradation throughout the tissue (*2*). The source of morphogen production is a key determinant of the gradient shape in that the gradient amplitude depends on the morphogen flux through the source boundary. However, in many cases the formation of a morphogen source is a dynamic process that depends on ongoing source specification as well as tissue growth. How the dynamics of morphogen source formation contributes to the formation of the morphogen gradient itself is poorly understood. Here we address this question by developing a mathematical model of a dynamic morphogen source in a growing tissue inspired by Shh gradient formation in the developing vertebrate neural tube.

In the neural tube, a morphogen gradient of Shh forms along the dorsoventral (DV) axis in the ventral to dorsal direction. Shh is produced by two distinct sources - the notochord and the floor plate (Fig. 1A). Shh expression first occurs within the notochord, a rod-like organ which is present from the onset of neurulation and is positioned underneath the neural tube (*3*). Prior to posterior neural tube closure, neural plate cells are competent to differentiate into floor plate, a specialized group of cells at the ventral midline (Fig. 1A) that directs spinal cord patterning and axon guidance (*4–7*). Early in vitro experiments have shown that floor plate specification can be induced by signals derived from either notochord or floor plate (*4*). In amniotes, the notochord is required for posterior floor plate formation (*7–9*) and Shh has been identified as the notochord-derived signal that mediates floor plate induction (reviewed in (*10*)). Shh secreted from the notochord spreads to the neural tube, where it forms an exponential gradient (*11, 12*). Over time, the decay length of the Shh gradient in the mouse neural tube remains approximately constant, while its amplitude increases several fold in the course of three days (*12*). Besides floor plate, Shh controls the formation of an organized pattern of ventral neural progenitor subtypes along the DV axis (*13*). In this process, which takes place predominantly in the first 24h of spinal cord development in mouse and chick, Shh signaling influences the dynamics of an underlying transcriptional network that specifies distinct molecular identities at different positions (*5, 14–18*).

**Fig. 1.**
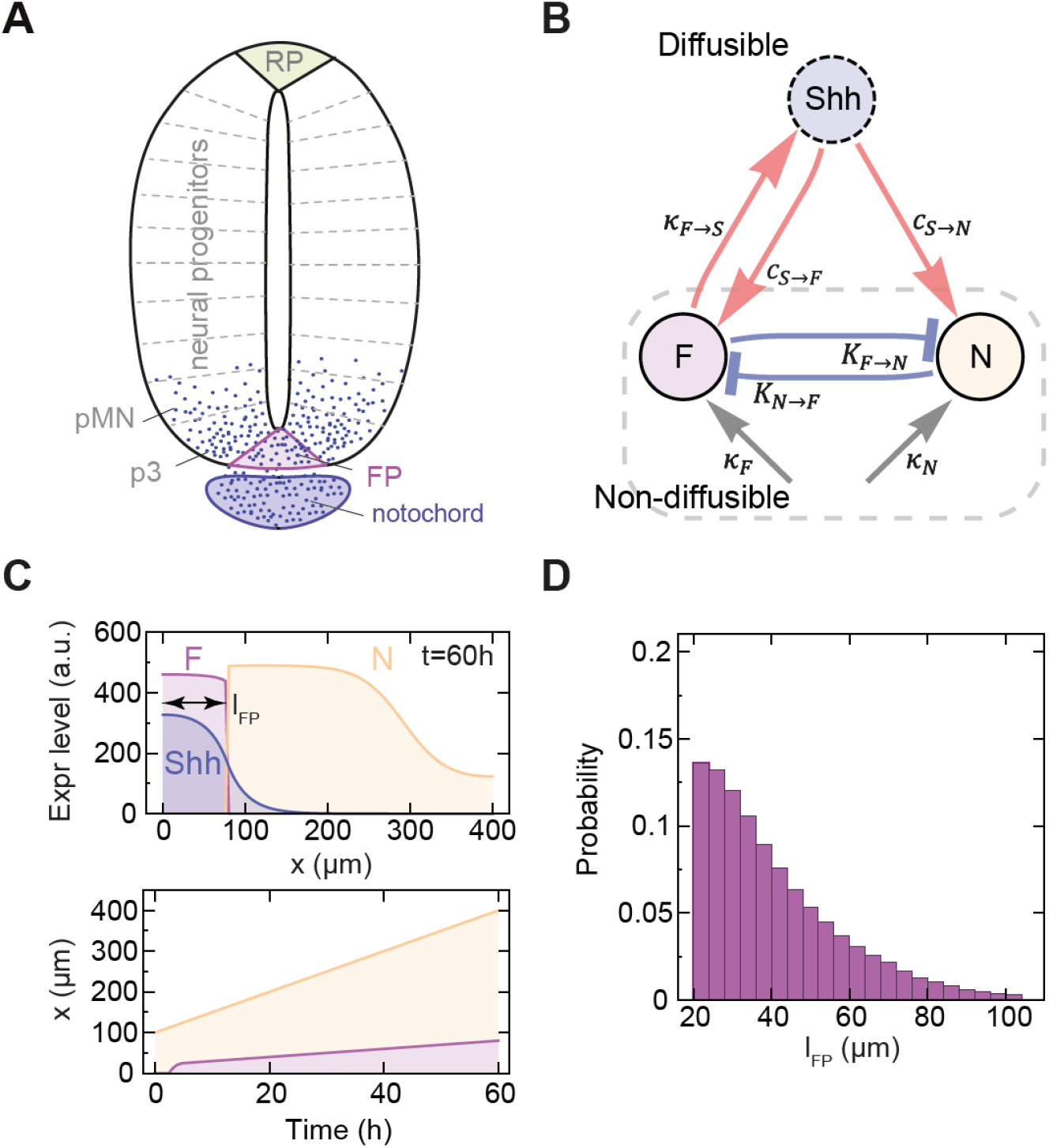
Model of floor plate formation in a growing tissue. **A**. Schematic illustration of the neural tube with indicated notochord, floor plate (FP), neural progenitor domains including p3 and pMN (grey), and roof plate (RP). Shh is depicted with blue dots. **B**. Schematic of the interactions in the model of FP formation considered in this study (Eq. 1). The nodes correspond to the non-diffusible transcription factors F and N, which define floor plate (F; purple) and neural progenitor identities (N; yellow), and to diffusible Shh morphogen (blue). Edges indicate activation strengths (red), repressor binding affinities (blue), and uniform activation (grey). **C**. Spatial pattern of FP and neural progenitor domains at the end of the simulation (top) and over time as the tissue grows (bottom). *l*_*FP*_ (double-arrow) indicates the FP size at *t* = 60h. **D**. Probability distribution of *l*_*FP*_ in successful solutions of the computational screen, n = 169 979.

While the spatiotemporal profiles of Shh levels, gene expression and growth in the mouse neural tube have been measured (reviewed in (*13*)), the mechanisms that underlie the formation of the Shh gradient and the specification of the floor plate are still poorly understood. One question is how the size of the floor plate domain is determined. Because floor plate specification depends on Shh and the floor plate itself produces Shh, this creates the potential for positive feedback to transiently contribute to the ongoing changes of the floor plate domain (*4, 15*). Gene regulatory interactions also contribute to the specification of floor plate (FP) identity and are likely relevant for defining the boundaries of this domain. For instance, floor plate identity genes such as Arx are repressed by the transcription factor Nkx2.2 which is expressed in the adjacent p3 domain, and conversely FoxA2, an transcription factor initially expressed in the floor plate and p3 domain, represses Nkx2.2 and p3 identity (*5, 19, 20*). These interactions occur within a growing tissue, raising the question of how floor plate formation is affected by tissue growth. A second question is how the profile of the Shh morphogen gradient depends on the formation of the floor plate. While the Shh amplitude increases over time, it is unclear what is the differential contribution of the notochord versus floor plate to the gradient shape.

In order to understand the contribution of different factors to the size of the floor plate and the Shh morphogen gradient dynamics, we developed a theoretical model supported by experimental evidence. The model is based on a simplified description of the interactions between fate determinants specific to the floor plate, the adjacent neural progenitor domains and Shh, coupled to a reaction-diffusion equation describing Shh spreading on a growing domain. By performing a parameter screen, we found that there are distinct possible mechanisms of floor plate formation. In one class of mechanisms, Shh produced within the floor plate is necessary for floor plate formation, while in another it is dispensable. The experimental evidence supports the latter mechanism. In this class, the floor plate size depends on different factors at distinct times. Initially, the floor plate domain is rapidly established is response to Shh and depends critically on the strengths of gene regulatory interactions. Subsequently, the size of the floor plate is passively expanded by tissue growth, leading to scaling of the floor plate with the tissue length. This growth of the floor plate, together with continuous Shh flux from the notochord, contribute to the increasing Shh gradient amplitude over time.

## Methods

### Model

In order to capture the dynamics of floor plate formation and its relationship with the Shh morphogen gradient within the growing neural tube, we developed a reaction-diffusion model that integrates a thermodynamic description of relevant gene interactions. The model represents a simplified interaction network between three species: F, N and Shh (Fig. 1B). F and N represent non-diffusible factors that are linked through effective cross-repressive interactions and define the identity of floor plate or adjacent neural progenitor domains, respectively. Shh represents the diffusible ligand which activates both N and F. The system dynamics is described by:

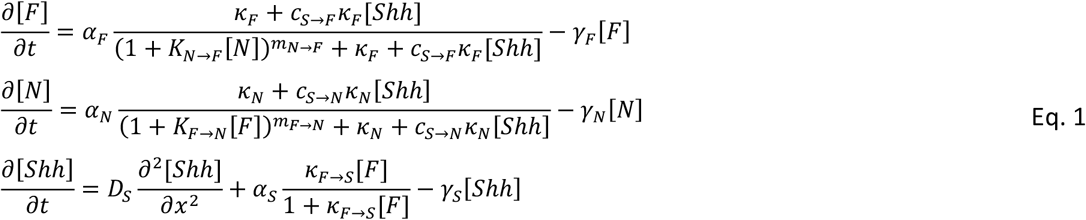

where [*F*], [*N*] and [*Shh*] are the concentrations of the interacting species, *D*_*S*_ is the diffusion constant of Shh, *κ*_*F*_, *κ*_*N*_, are uniform activation constants, *K*_*N*→*F*_, *K*_*F*→*N*_ are repressor binding affinities, *c*_*S*→*F*_, *c*_*S*→*N*_, are morphogen activation strengths relative to uniform activation, and *κ*_*F*→*S*_ is the activation strength of Shh ligand production by the floor plate. Note that Shh is not produced in the floor plate if either [*F*] or *κ*_*F*→*S*_ = 0. To reduce the number of free parameters, we set the production rates *α* to 0.1 h^-1^, degradation rates *γ* to 0.72 h^-1^ (= 2⋅10^−4^ s^-1^), and *D*_*S*_ = 0.11 μm^2^ s^-1^ – these values are of similar order to the values measured for other morphogens (reviewed in (*21*)) and to inferred values for Shh in other studies (*22–24*). The exponents that quantify nonlinearity *m*_*N*→*F*_, *m*_*F*→*N*_ were set to 3. All gene expression levels and morphogen concentration depend on position 0 ≤ *x* ≤ *L*, and time *t* ≤ *t*_*end*_.

We model a one-dimensional tissue that grows linearly, starting from an initial length of *L*_0_ = 100 μm with a default growth rate *k*_*p*_ = 5 μm/h. Thus, the tissue reaches the default final length *L*_*end*_ = 400 μm at time *t*_*end*_ = 60 h, which is similar to the experimentally measured DV length of the neural tube after 60 hours of development (*17*). In order to study how the tissue growth rate affects the model behaviour, we set the final tissue length such that it corresponds to growth rates from *k*_*p*_ = 0 μm/h (*L*_*end*_ = 100 μm, non-growing condition) to *k*_*p*_ = 50 μm/h (*L*_*end*_ = 3100 μm). Simulations are performed on a growing one-dimensional tissue divided into 100 spatial bins that mimic the discrete cellular structure of the tissue. Tissue growth is implemented by increasing the size of bins, such that both dilution and advection effects resulting from growth are handled by the numerical integration scheme. The integration scheme uses first order finite differences of the spatial derivatives, whilst time steps are handled using Heun’s scheme, which is a second order method. The F and N domains are defined according to the gene expression levels, such that [*F*] > [*N*] defines the F domain, and [*F*] < [*N*] the N domain. Throughout the text, we refer to the F domain also as FP or FP domain.

The initial conditions of the model are such that at *t*=0 N is expressed uniformly across the tissue ([*N*]_*init*_ = 10 a.u. for 0 ≤ *x* ≤ *L*_*end*_), reflecting the transient expression of N in future FP cells, while there is no initial expression of *F* (i.e. [*F*]_*init*_ = 0) (*5*). To represent Shh secreted by the notochord (*25–27*), we consider different initial conditions, including a transient burst of Shh at position *x* = 0 within the tissue ([*S*]_*init*_ = 100 a.u. at *x* = 0, and [*S*]_*init*_ = 0 for 0 < *x* ≤ *L*_*end*_), or a constant flux of Shh *j*_*Shh*_ from the ventral boundary of the neural tube at 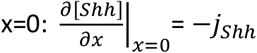. The *j*_*Shh*_ can be abruptly removed at a specific time *t*_*off*_.

### Computational screen

A priori, the model in Eq. 1 (Fig. 1B) may lead to no stable formation of a F domain, or have a F domain that extends across the entire tissue. In order to identify parameter sets that result in biologically plausible FP formation (Fig. 1C), we define the following constraints: (i) the emerging pattern has two domains, F and N, with the F domain starting at the ventral end (*x* = 0), (ii) the Shh profile decays monotonically as a function of position x, (we achieve this by imposing an upper bound on Shh concentration at *L*_*end*_ of 2% of [*S*]_*init*_, hence avoiding rare cases in which the FP forms at the dorsal end), (iii) [*Shh*] and [*F*] concentrations differ at most by an order of magnitude, that is 0.1 ≤ [*Shh*]/[*F*] ≤ 10; this is a technical assumption to keep the relative range of [*Shh*] and [*F*] bounded, as effectively *κ*_*F*→*S*_ acts as scaling factor for [*F*] (see Eq. 1) and *κ*_*F*→*S*_ is varied by 6 orders of magnitude (see below). We have two additional criteria that narrow down the solutions to biologically realistic spatial and temporal scales: (iv) the length of the F domain at *t*_*end*_ is between 5% and 25% of tissue length (consistent with 7.5% of tissue length measured at *t* = 60 h in(*17*)); this restricts the space of successful solutions to F lengths from 20 μm to 100 μm at *t*_*end*_, (v) FP is established between 2.5h and 20h; the time of FP establishment *T*_*est*_ is defined as the time at which [*F*]>[*N*]. The minimal *T*_*est*_ of 2.5 h avoids oversampling the parameter space in the region of establishment times on the order of minutes, which are inconsistent with experimental results (*17*).

The computational search of parameter space is performed by random walk in the logarithmic parameter space. The following parameters are varied over 6 orders of magnitude, *c*_*S*→*F*_, *c*_*S*→*N*_, *κ*_*F*→*S*_, *K*_*N*→*F*_, *K*_*F*→*N*_ from 0.005 to 5000, and *κ*_*F*_, *κ*_*N*_ from 5⋅10^−6^ and 5. The initial parameter set is selected randomly. If this set is not fulfilling all success criteria (i)-(v), another random set of parameters is selected. If the selected parameter set satisfies all success criteria, the next set of parameters is generated based on the preceding set, by multiplying screen parameters by random numbers from log-normal distribution with 0 mean and 0.2 standard deviation. This multiplication by random numbers is performed iteratively until predefined number of parameter sets is visited. The computational search was started independently 10 times with each search visiting 40 000 parameter sets, and the sets of identified successful solutions were combined.

### Mouse lines

All work with animals was approved under the license BMWFW-66.018/0006-WF/V/3b/2016 from the Austrian Bundesministerium für Wissenschaft, Forschung und Wirtschaft. All procedures were performed in accordance with the relevant regulations. The following mouse lines were previously described: Sox2^CreERT2^ (JAX:017593, (*28*)), Shh^CreERT2^ (JAX:005623, (*29*)), and Shh^Flox^ (JAX:004293, (*30*)). Transgenic strains were maintained on a CD-1 background. Sox2^CreERT2/+^ was crossed to Shh^Flox/+^ to generate Sox2^CreERT2/+^, Shh^Flox/+^ mice, which were further crossed to Shh^Flox/+^ to generate Sox2^CreERT2/+^, Shh^Flox/Flox^ embryos. Females were injected at E7.5 with 3mg tamoxifen, and embryos were collected at E10.5. Shh^CreERT2/+^ mice were crossed to Shh^Flox/+^ mice to generate Shh^CreERT2/Flox^ embryos. Females were injected at either E5.5 and E6.5 with 3 mg tamoxifen each, or at E8.5 with 3 mg tamoxifen; and embryos were subsequently collected at E10.5.

### Immunohistochemistry

Embryos were fixed in PBS with 4% PFA for 2 hours, embedded in gelatin, and cryosectioned. Transverse brachial sections were pre-blocked in PBS with 0.1% Triton X-100 (PBST), 1% BSA for 1 hour, incubated overnight at 4°C in PBST with the following primary antibodies: mouse SHH supernatant (DSHB, 1:70) and sheep ARX (R&D systems, 1:100), washed in PBST, incubated with fluorescently labelled secondary antibodies (Jackson ImmunoResearch, donkey anti mouse IgG-Cy3, 1:1000; donkey anti sheep IgG-FITC 1:250) and DAPI (ThermoFisher, 1:100) in PBST for 2h at room temperature, washed in PBST and mounted for imaging.

### Imaging and quantification of mouse sections

Immunostained sections were imaged using a ZEISS LSM800 Axio Observer Z1 confocal microscope with a Plan-Apochromat 40x/1.2 water objective, at 0.8x zoom, with a Z-stack composed of 8 Z-planes 0.6 μm apart. Subsequent image analysis was performed using Fiji (macro modified from (*31*), (*32*)). After maximum intensity projection, the ARX-positive floor plate area was quantified using the “Threshold” function in default mode. To quantify the SHH fluorescence intensity (FI), a 14μm-wide ROI was aligned along the apical surface of the neural tube, starting from the ventral midline, to obtain a mean pixel intensity calculated across the 14μm ROI width as a function of ventral-to-dorsal position. The SHH profiles were subsequently analyzed in Python. For every profile, background, defined as the minimum FI within 10-90% of DV length, was subtracted and the FI was smoothed with a moving average filter (window size of 5μm). The mean intensity profile was calculated for each genotype. Then, x=0 was determined by finding the position of the two SHH intensity peaks for the mean profile (located at the ventral and dorsal edge of the floor plate respectively) by calculating local maxima, and setting the position of the peak corresponding to the dorsal edge of the floor plate as x=0 for all profiles. x=0 for Sox2^CreERT2/+^, Shh^Fl/Fl^ and Shh^CreERT2/Fl^ profiles was set to the same position as in their respective control mean profiles. SHH background subtraction was also performed on the mean profiles. SHH profiles were normalized to the mean maximum intensity of the control group.

## Results

### Floor plate formation independent of floor plate-derived Shh relies on strong network interactions

In order to understand how the FP size is determined, we performed a computational screen to identify parameter sets that lead to FP formation using a reaction-diffusion model that incorporates genetic interactions that influence the specification of FP identity and tissue growth (Fig. 1B), as described above. This resulted in the identification of 169 979 successful parameter sets out of 400 000 visited parameter sets at the end of the simulation at *t* = 60h (Fig. 1C). Within this set of successful solutions, FP sizes followed a half-normal distribution, such that FP sizes between 20μm and 40μm accounted for 66% of successful solutions, while 4% had FP size ≥ 80μm (Fig. 1D).

To understand how FP size depends on model parameters, we examined the parameter distributions for subsets of networks that produced FP sizes within a defined range. The distributions of model parameters for different sized FPs showed that the model can produce all possible FP sizes over broad parameter ranges (Fig. S2A). In particular, contrary to our expectation that the strength of activation of F by Shh should influence FP size, the distribution of *c*_*S*→*F*_ did not vary with FP size (Fig. S2A). This suggests that FP size is indirectly affected by other model parameters. To understand the sensitivity of FP size to model parameters, we performed sensitivity analysis focusing on a subset of networks that yield floor plates with a relative size of 20% of DV length. We measured how the relative size of the FP changed upon 10 fold up- and down- regulation of each parameter relative to its mean value for that subset, while all other parameter values were held constant. The sensitivity analysis revealed that FP size was highly sensitive to changes in all parameters, except *κ*_*F*→*S*_, for which the dependence was weak (Fig. 2A). In particular, FP size increased when F was strongly activated and N was deactivated or repressed. Conversely, FP size decreased when N was strongly activated and F was deactivated or repressed. Altogether, this indicated that FP size is influenced by all interactions within the network, but is least sensitive to Shh production in the FP. This suggests that the FP may form in a manner that is largely independent of the Shh that it produces.

**Fig. 2.**
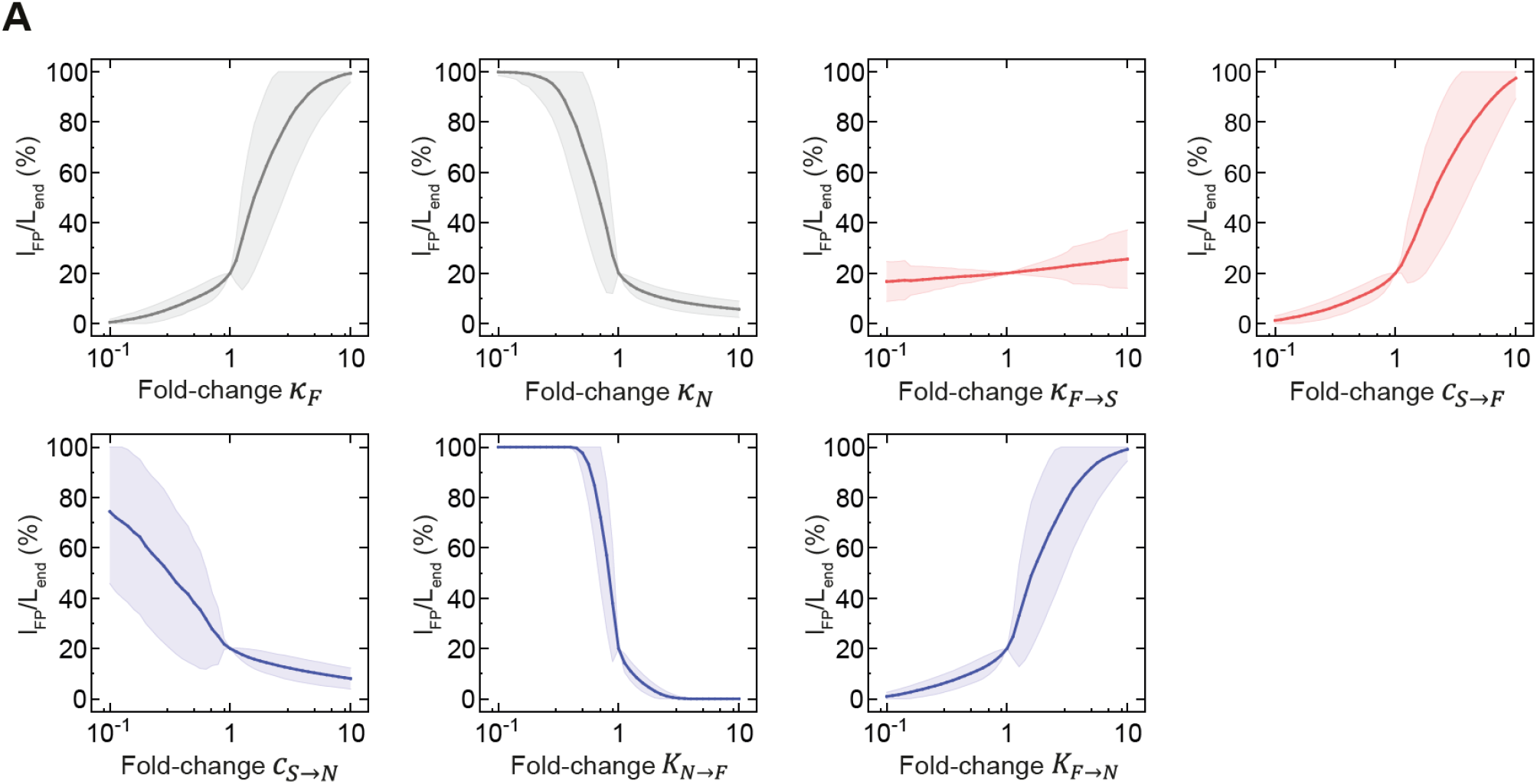
FP size sensitivity to perturbations of model parameters. **B**. Mean relative FP size at the end of the simulation upon perturbation of the indicated model parameter for a subset of 214 randomly selected networks with relative FP size of 20%. The parameters are modified in 40 equally distributed logarithmic steps from 0.1 to 10-fold of their value. The shaded regions are SE.

To further investigate how Shh production in the FP affects FP size, we set the production term *κ*_*F*→*S*_ to 0, which corresponds to a case in which there is no floor plate-derived Shh (Shh^FP^). We then compared the full model to the perturbed case by quantifying the change in FP size: 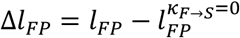, where 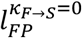 is FP size in the perturbed case. In most cases, the perturbed networks exhibited one of three types of responses, in which the FP size was either maintained, decreased, or completely lost (Fig. 3A). This indicates that the FP in these networks was insensitive, partially sensitive or completely sensitive to its own production of Shh, respectively. In a small fraction corresponding to ∼1.5% of all cases, the FP size increased when the production term *κ*_*F*→*S*_ was perturbed (Fig. 3A). This could result from a situation in which Shh produced in the FP is required to maintain N expression, thereby restricting the expansion of FP dorsally. Together, this indicates that depending on the value of other parameters, the magnitude of *κ*_*F*→*S*_ can significantly affect FP size.

**Fig. 3.**
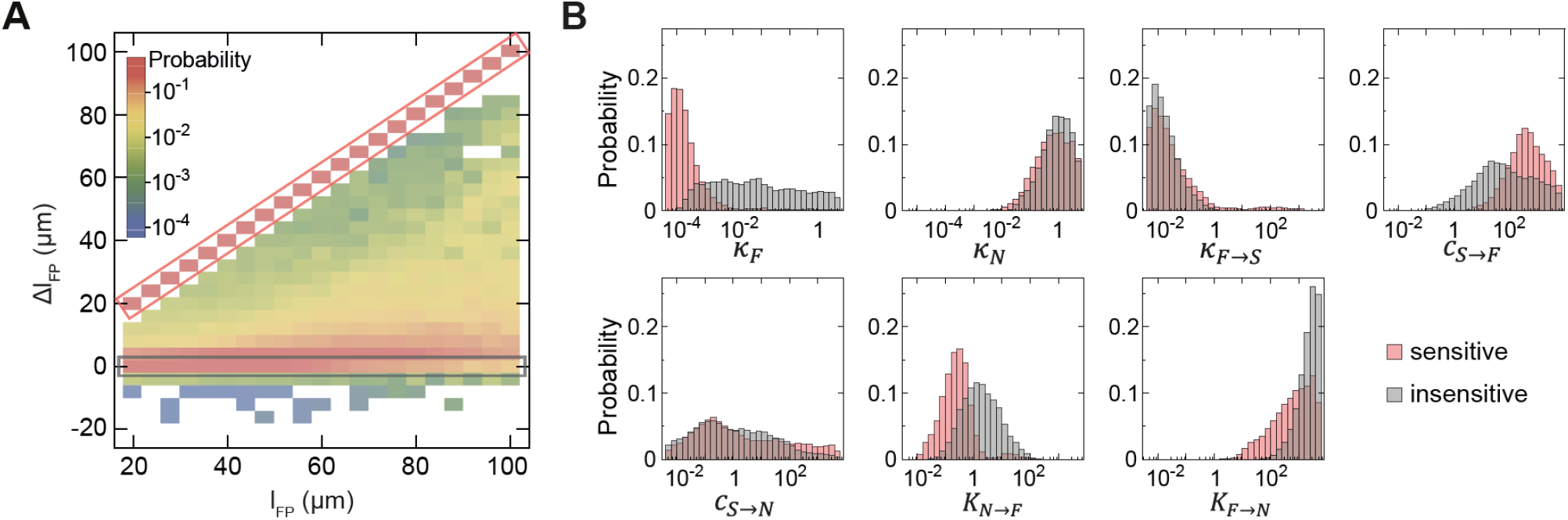
FP formation can occur by different mechanisms, dependent or independent of FP-derived Shh. **A**. Probability distribution of the change in FP size (Δ*l*_*Fp*_) for successful solutions with production of Shh by FP switched off (*κ*_*F*→*S*_ = 0) compared to the default model (*κ*_*F*→*S*_ > 0). Rectangular frames indicate solutions with FP formation dependent on Shh^FP^ (Δ*l*_*FP*_ = *l*_*FP*_; sensitive; red) and independent of Shh^FP^ (Δ*l*_*FP*_= 0; insensitive, grey). Colours correspond to log-scaled conditional probability of observing Δ*l*_*Fp*_ for a given *l*_*FP*_(legend), n = 168 074 (all solutions except 1905 with FP extending across the whole tissue at *t* = 60h). **B**. Distribution of model parameters for Shh^FP^-sensitive (red) and insensitive (grey) classes of solutions, sensitive (n = 45 336), insensitive (n = 66 827).

To understand what distinguishes these classes of networks, we compared the parameter distributions within each class (Fig. 3B, Fig. S1). The most notable difference was that Shh^FP^-sensitive solutions required smaller basal activation *κ*_*F*_ and at the same time had overall higher strength of activation of F by Shh (*c*_*S*→*F*_), consistent with their strong dependence on Shh. By contrast, solutions that are insensitive to Shh^FP^ require strong basal activation and have lower *c*_*S*→*F*_. Furthermore, insensitive networks have higher repression of F by N. This suggests that in the insensitive class, FP formation results from strong basal uniform activation and repression by Shh-dependent repressors, with weak initial activation by Shh and with no activation by FP-derived Shh.

### Rapid FP formation is followed by expansion via tissue growth

To investigate how FP formation can occur without being affected by Shh production in the FP, we analysed the time course of FP formation in the insensitive class of solutions compared to the sensitive class. We observed that as the overall tissue length increases over time, the FP size also increases (Fig. 4A, B). Notably, the FP size occupies a constant fraction of the overall tissue length from 10 h onwards (Fig. 4A’, B’). This indicates that after this time point, the FP size scales with tissue growth (Fig. 4A’, B’). These dynamics were unchanged in the absence of Shh^FP^ in the insensitive class of solutions (Fig. 4C, C’), while in the sensitive class, the FP domain was lost after 10 h (Fig. 4D, D’).

**Fig. 4.**
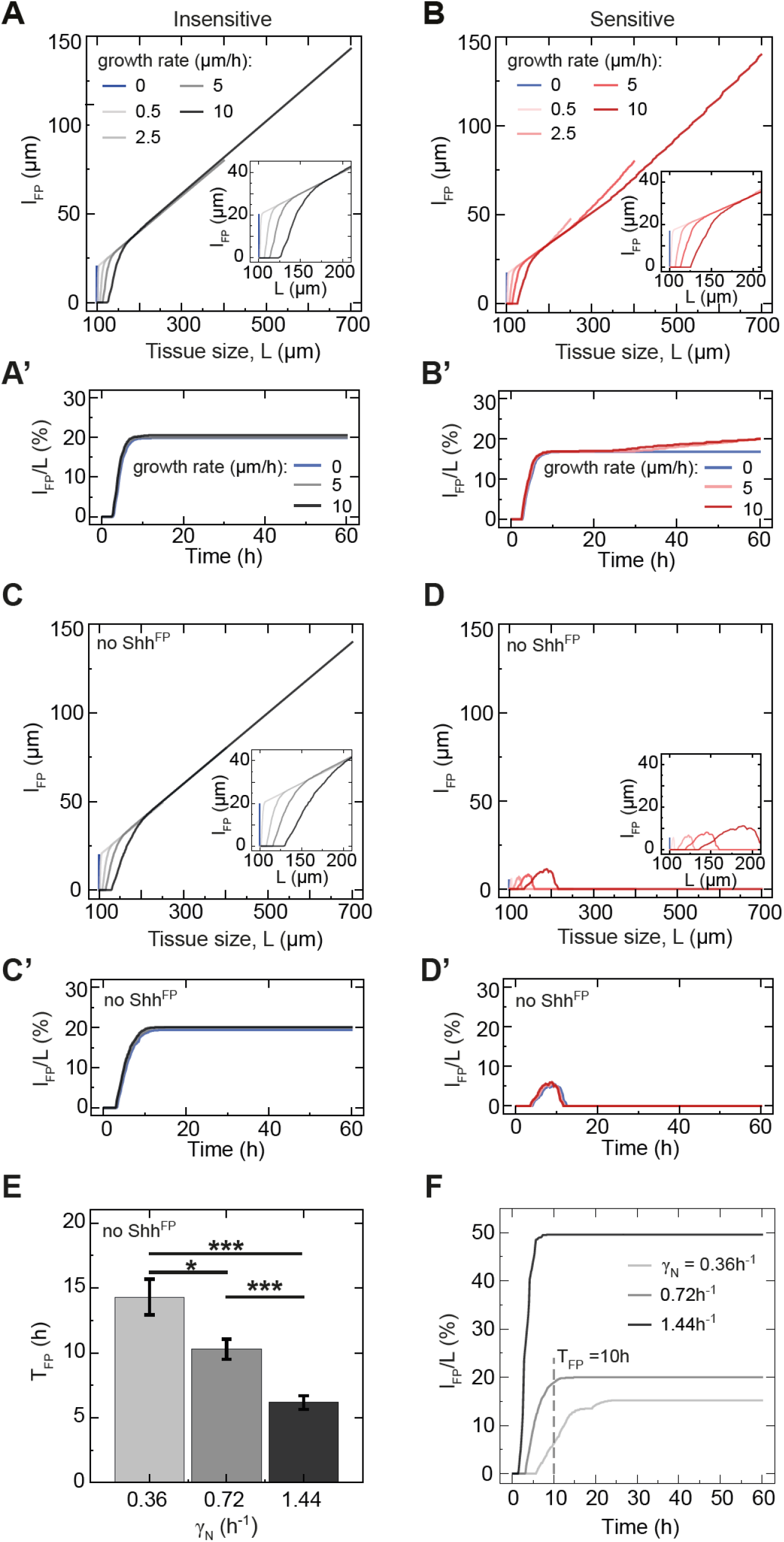
Two time scales of FP formation. **A, B**. Absolute FP size as a function of the increasing tissue size L over time for Shh^FP^-insensitive (A) and sensitive (B) solutions for the default model with Shh^FP^. The tissue growth rate is varied from *k*_*p*_=0 μm/h to 10 μm/h (color code in the legend). The inset shows a magnified view for small tissue sizes. **A’, B’**. Relative FP size as a function of time for insensitive (A’) and sensitive (B’) solutions with Shh^FP^ and varied growth rate. Growth rates *k*_*p*_=0 μm/h (blue), 5 μm/h (grey/pink), and 10 μm/h (darker grey/red) are shown. **C, D**. The same as A and B, respectively, but in the absence of Shh^FP^. **C’, D’**. The same as A’, B’ respectively, but in the absence of Shh^FP^. A-D’, each curve is the mean of n=10 solutions. **E**. Average FP formation time *T*_*FP*_ (defined as the time at which FP reaches its final relative size) as a function of the degradation rate of N (*γ*_*N*_), see Eq. 1. Pairwise comparisons two-tailed *t*-test: * 0.05 ≥ *P* > 0.01; *** 0.001 ≥ *P* > 0.0001. n=10 per condition, error bars SEM. **F**. Relative FP size as a function of time for different *γ*_*N*_. The default condition is *γ*_*N*_= 0.72h^-1^, dashed line indicates the position of the average *T*_*FP*_ = 10 h for that condition.

The scaling of the FP with tissue size from 10 h onwards implies that growth passively expands the FP domain after this time. To test this, we varied the growth rate *k*_*p*_ from 0.5 μm/h to 10 μm/h, resulting in tissues with different final sizes at 60h, from 100μm to 700μm, respectively. Strikingly, the relative FP size was growth rate-invariant, indicating that the FP scaled with the tissue sizes produced in different growth regimes (Fig 4A’, B’). Weak deviations from perfect scaling were only observed for large FPs from the sensitive class of networks (Fig. 4B’).

The time course analysis suggests that there are two time scales of FP formation (Fig. 4A’, B’). Initially, the FP rapidly, within 10 h, reaches a size that is defined by the model parameters. Subsequently, the FP scales with tissue size, hence the absolute FP size is defined by the amount of growth. Consistent with this, in the absence of growth, FP formation is completed within 10h (Fig. 4A, B). We then asked which parameters control this initial rapid phase of FP formation. While most of the parameters in the computational screen do not influence the temporal dynamics (Fig. S3), the time it takes for the relative FP size to become constant (*T*_*Fp*_) was influenced by the degradation rate of the reacting species, notably *γ*_*N*_ (Fig. 4E). A 2-fold increase in the degradation rate of N leads to an approximately 2-fold decrease in *T*_*Fp*_ (Fig. 4E, F).

The scaling of FP with tissue size after 10 h suggests that the FP is established in response to Shh initially present in the system and after 10 h is not sensitive to changes in Shh concentration. To test whether this is the case, we investigated how the FP size changes when the Shh input is configured differently. In our model so far, Shh is initially provided as a fixed pulse of 100 a.u. in the first spatial bin at *t*=0. In the absence of this initial pulse, FP formation does not occur. However, in the embryo, Shh is continuously produced by the notochord. To simulate the effect of the notochord, we compared our default model with an initial pulse of Shh to a model with a constant flux of Shh, representing the notochord. We found that for a specific magnitude of flux, the model produced a similar relative FP size as in the model with a pulse (Fig. 5A). However, increasing flux lead to an approximately linear increase in relative FP size. This was the case for both sensitive and insensitive classes of solutions (Fig. 5A, B). Crucially, the presence of flux within a test range spanning >4 orders of magnitude did not alter the temporal dynamics of FP formation – FP size still scaled with tissue size after 10 h (Fig. 5C). This supports the idea that any changes in the flux contribute to FP formation only early on, but that the FP does not respond to the continuous flux of Shh after this time point.

**Fig. 5.**
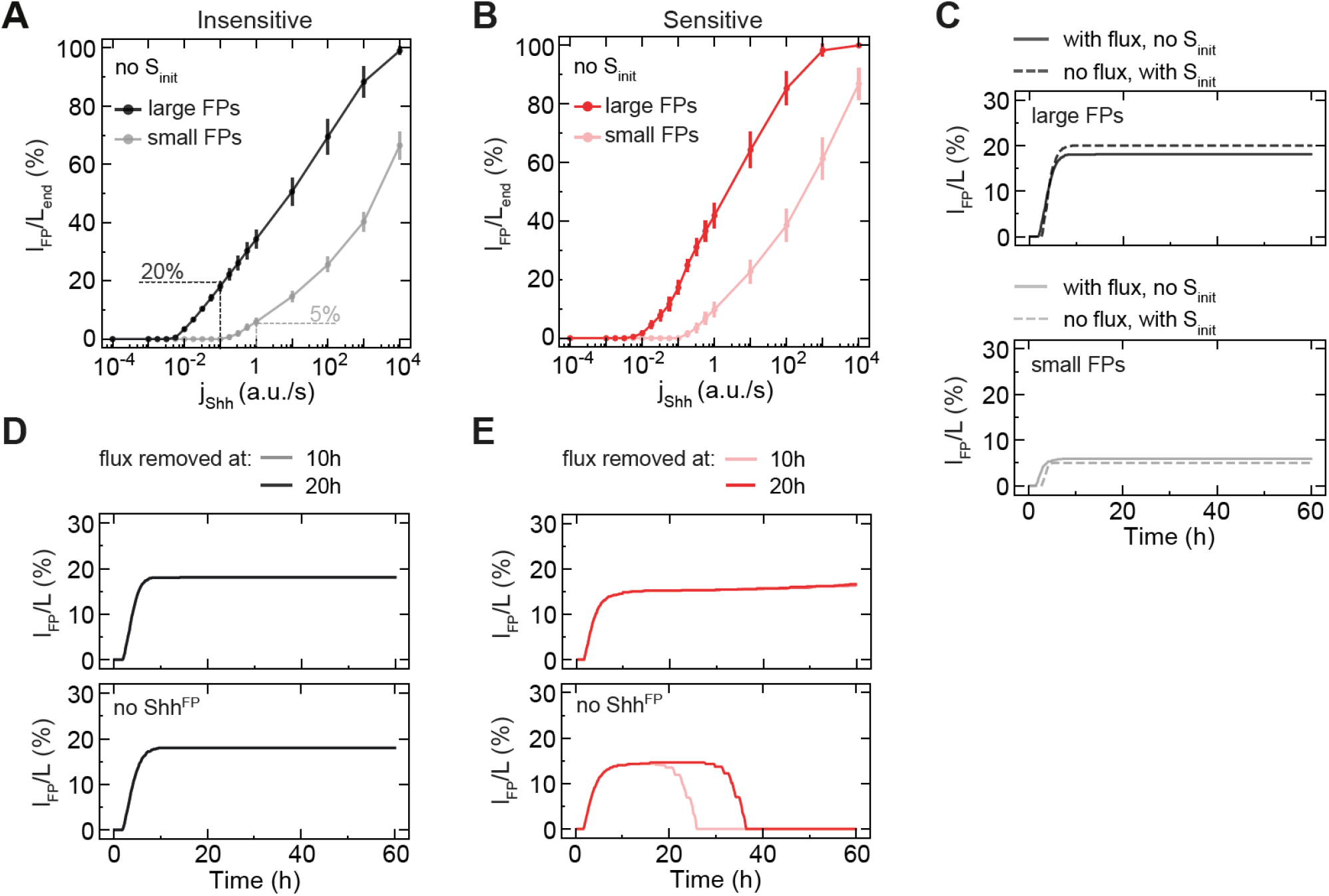
The temporal dynamics of FP formation is independent of continuous Shh flux. **A, B**. Relative FP size for insensitive (A) and sensitive (B) solutions with varied flux of Shh and without initial Shh. Solutions resulting in large FPs (*l*_*Fp*_ /*L*_*end*_=20%; darker grey/red) and small FPs (*l*_*Fp*_ /*L*_*end*_=5%; lighter grey/red) for default conditions (no flux, but with initial Shh) are shown separately. The magnitude of flux that yields approximately the same FP size as the default condition (no flux, but with initial Shh) is indicated with dashed lines. **C**. Relative FP size as a function of time for default condition (dashed) vs analogous flux condition (solid). *j*_*Shh*_=0.1 a.u./s for large FPs (top) and *j*_*Shh*_=1 a.u./s for small FPs (bottom). **D, E**. Relative FP size with flux abruptly removed at the indicated time (colored) for insensitive (D) and sensitive (E) solutions for large FPs. With Shh^FP^ (top), without Shh^FP^ (*κ*_*F*→*S*_ = 0; bottom). A-E, the average relative FP size at 60h is shown, n=10 per point. A and C, error bars SEM.

To test this directly, we performed a simulation in which there was no initial pulse of Shh and instead, Shh flux was provided for 10 or 20 hours and subsequently removed. This showed that both in the sensitive and insensitive classes of networks, external Shh input is required for only 10h, and subsequently the FP is maintained without the need for continuous Shh flux (Fig. 5D, E). Notably, in the absence of Shh production in the FP, the FP was maintained after flux removal in the insensitive class of networks, but was lost in the sensitive class (Fig. 5C, E). This indicates that in the sensitive class, FP maintenance requires either continuous flux or Shh^FP^, while in the insensitive class, neither of these sources of Shh production is required for FP maintenance. Thus, this indicates that the insensitive class of networks is bistable with respect to Shh.

Altogether, these results show that in the subset of Shh^FP^ insensitive networks, the FP forms independent of Shh^FP^ because of the fast time scale of initial FP formation, which is set by the degradation rate of N. The FP is maintained independent of Shh due to the hysteresis inherent to this dynamical system with respect to the Shh input.

### FP amplifies the Shh gradient amplitude at late stages

Our model indicates that the initial FP formation depends on the Shh gradient, however, the contribution of different factors to Shh gradient formation is unclear. To address this, we analyzed the Shh gradient shape and asked how it depends on different conditions and model parameters. The decay length *λ* of steady state exponential gradients formed by diffusion with diffusion coefficient D, uniform degradation with rate k and localized production is equal to 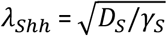 (*33*). Thus, because the diffusion and degradation parameters of Shh in our model are fixed, the decay length of the Shh gradient in the receiving tissue is expected to reach a constant value, corresponding to 23.45 μm. Consistent with this, we find that at *t* = 60 h, *λ*_*Shh*_ estimated from a fit to the exponentially decaying part of the Shh gradient in the set of successful solutions has a narrow distribution centered at 23.32 ± 0.09 μm (mean ± SE) (Fig. S4A). By contrast, the Shh amplitudes of successful solutions exhibited a broad ordinary normal distribution in the range between 100 and 500 a.u. with ⟨*A*_*Shh*_⟩ = 293 ± 75 a.u. (mean ± SE) (Fig. S4B). Therefore, the Shh gradient changes mainly through its amplitude.

To understand what factors influence the Shh amplitude, we first asked how it depends on changes of the model parameters. We found that the Shh amplitude and the FP size varied in a qualitatively similar manner with most parameter values (Fig. 6A compare to Fig. 2B). There was one notable exception – while the FP size did not depend strongly on the strength of Shh production by the FP, *κ*_*F*→*S*_, the Shh amplitude increased with increasing *κ*_*F*→*S*_. Thus, while Shh^FP^ is dispensable for regulating the FP size, it influences the Shh gradient amplitude. This raises the possibility that changes in the amplitude of Shh over time results from changes in the net production of Shh^FP^ over time.

**Fig. 6.**
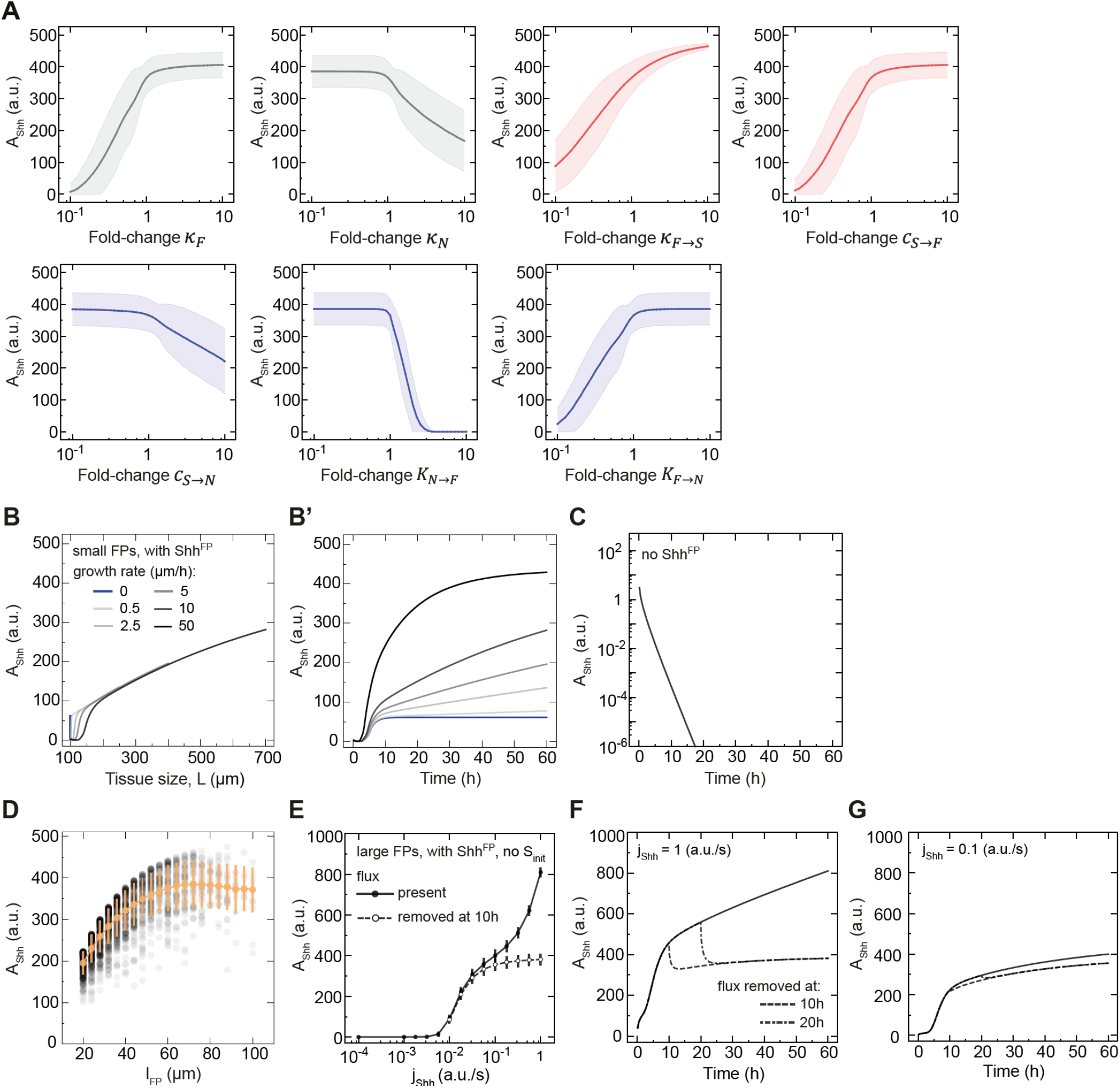
The increase in Shh amplitude over time depends on floor plate growth. **A**. Sensitivity of the Shh amplitude at the end of simulations to parameter perturbations. The parameters are changed in 40 equally distributed logarithmic steps from 0.1 to 10-fold of their value. Networks with *l*_*Fp*_ /*L*_*end*_=20% were randomly selected (n = 214, including n=100 sensitive, n=100 insensitive and n=14 partially sensitive). The shaded regions are SE. **B, B’**. Shh amplitude as a function of tissue size (B) and time (B’) for insensitive solutions with Shh^FP^. Growth rates from *k*_*p*_=0 μm/h to 50 μm/h are color-coded, n=10 per condition, sampled every 10 min. **C**. Shh amplitude as a function of time in the absence of Shh^FP^. The curves are identical for all n=10. **D**. Shh amplitude as a function of FP size. The yellow points correspond to mean *A*_*Shh*_ for a given *l*_*Fp*_, the number of samples per point from n=23 116 (*l*_*Fp*_ =20μm) to n=497 (*l*_*Fp*_ =100μm), error bars SE. Randomly selected points from successful solutions (black), n=3000. **E**. Shh amplitude for large FPs with varied flux of Shh and no initial pulse of Shh. The notochord flux is present at all times (black solid) or is removed at *t*_*off*_=10h (white dashed). **F**. Shh amplitude over time for large FPs with flux removed at specific times. The flux *j*_*Shh*_=1 a.u./s is present (solid) or removed at 10h (dashed), or at 20h (dot-dashed). **G**. The same as F, but for 10-fold lower flux *j*_*Shh*_=0.1 a.u./s. E-G, n=10 per condition.

To test this, we analysed the temporal profiles of Shh gradient formation. Similar to the FP size, which continuously increases over time, the Shh amplitude also increases (Fig. 6B, B’). Furthermore, we observed two clear time scales of Shh amplitude change. Up to ∼10h after the simulation onset, the Shh amplitude increased rapidly. This was followed by a second phase, in which the increase was slower (Fig. 6B’). To test whether the increase in Shh amplitude over time is due to Shh^FP^, we compared the temporal dynamics in the default case to a situation in which the floor plate does not produce Shh (*κ*_*F*→*S*_ = 0). We found that for *κ*_*F*→*S*_ = 0 the Shh amplitude did not increase, but continuously declined and reached negligibly small values (<10^−4^ a.u.) after 10h (Fig. 6C). This supports the conclusion that the increase of Shh amplitude over time depends on Shh produced by the FP.

Dependence of the Shh amplitude on Shh^FP^ implies that the Shh amplitude should depend on the tissue growth rate in a similar manner to the FP. To test this prediction, we analysed the Shh amplitude and decay length over time and at different tissue growth rates. In the first 10h, the Shh decay length exhibited transient dynamics corresponding to a shift from the initial Shh present in the system to the decay length predicted by the fixed Shh diffusion and degradation rate 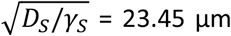 (Fig. S4C). After 10h, the decay length remained approximately constant for a wide range of growth rates (Fig. S4C). By contrast, the Shh amplitude increased with increasing growth rate (Fig. 6B, B’). Furthermore, the temporal changes in amplitude depended on the growth rate. In the absence of growth or at an unrealistically high growth rate of *k*_*p*_ = 50 μm/h, the amplitude reached saturation levels in less than 30h (Fig. 6B’). For low and intermediate growth rates, the Shh amplitude increased linearly over time between 10h and 60h (Fig. 6B’), as well as with respect to the growing tissue length L, indicating that the amplitude of Shh scales with the tissue size after 10h (Fig. 6B).

Together, our analysis suggests that tissue growth leads to an increase in the Shh amplitude by expanding the FP size and thus increasing Shh production (Fig. 6B). The size of the morphogen source, however, may be non-linearly related to morphogen flux through the source boundary and to the gradient amplitude, for instance due to the fact that newly produced molecules will be degraded before they spread to the source boundary at large source sizes (*34, 35*). Consistent with this, analysis of the relationship between FP size and Shh gradient showed that while there was substantial variation of Shh amplitudes for any specific FP size, on average the Shh amplitude varied non-linearly with respect to FP size (Fig. 6D). While small FP sizes, below ∼60 μm, were on average linearly related to the Shh amplitude, for large FPs the Shh amplitude saturates (Fig. 6D). This indicates that beyond FP size of ∼60 μm, the amplitude of Shh and the FP size vary independently.

Nevertheless, in the linear regime, the question remains of what the contribution of the floor plate versus continuous flux from the notochord is to the Shh amplitude dynamics. To investigate this, we asked how the presence of flux changes the dynamics of the Shh gradient amplitude, focusing first on the model without Shh production in the FP. We found that in the case with no initial pulse of Shh and no production of Shh by FP, increasing flux lead to increased Shh amplitude *A*_*Shh*_∼*j*_*Shh*_, as expected from a model of gradient formation by diffusion, degradation and localized production (*33*) (Fig. S5). Furthermore, in the presence of Shh production by FP (*κ*_*F*→*S*_>0), the increase in *A*_*Shh*_ occurred at smaller values of flux (Fig. 6E). This supports the conclusion that the presence of the FP amplifies the Shh amplitude. To understand the contribution of continuous influx of Shh from the notochord to the Shh profile dynamics, we turned off the flux at *t*_*off*_ = 10h. In the absence of Shh^FP^, this perturbation abrogated the formation of a Shh gradient. In the presence of Shh^FP^, the amplitude was reduced and the extent of this reduction depends on the magnitude of the flux (Fig. 6E). In a regime with very high flux (1 a.u./s, 10-fold higher than the default condition, cf. Fig. 5C), the flux has a strong contribution to the overall Shh production until the end of the simulation (53% ± 4% of *A*_*Shh*_, mean ± SEM, Fig. 6F), while for smaller flux values – the effect of continuous flux on the Shh amplitude was nearly undetectable (11% ± 9%, mean ± SEM, Fig. 6G). Thus, our analysis indicates that besides the floor plate, continuous flux from the notochord can potentially contribute to increasing the Shh gradient amplitude over time.

### Experimental validation of the model

Our model revealed that FP formation can in principle occur via different mechanisms, depending on whether FP-derived Shh contributes to FP formation. To determine which mechanism is relevant to the *in vivo* situation, we deleted Shh production from the floor plate, while leaving production in the notochord intact. To this end, we used embryos homozygous for a Shh^Flox^ allele and carrying one copy of the tamoxifen inducible Sox2^CreERT2^ (Methods), which is expressed in floor plate cells, but not in the notochord. Endogenous Shh expression in the mouse floor plate is initiated around E9.5 (*11, 36*), hence we injected mothers with tamoxifen at E7.5 of development to eliminate any Shh production in the floor plate. In this condition, at E10.5 of development, embryos had a significant reduction of approximately 95% in their Shh amplitude, while the floor plate size, assessed by Arx immunostaining, was unchanged compared to control littermates (Fig. 7A-C). This indicates that FP-derived Shh is not required for the formation of the floor plate. These results are consistent with prior Shh^FP^ deletion experiments, in which no effect was observed on FoxA2 expression (*14*), as well as with observations that the FP becomes refractory to Shh signaling from E9.5 onwards (*5*). Taken together, these results supports a Shh^FP^ insensitive mechanism of FP formation *in vivo*.

**Fig. 7.**
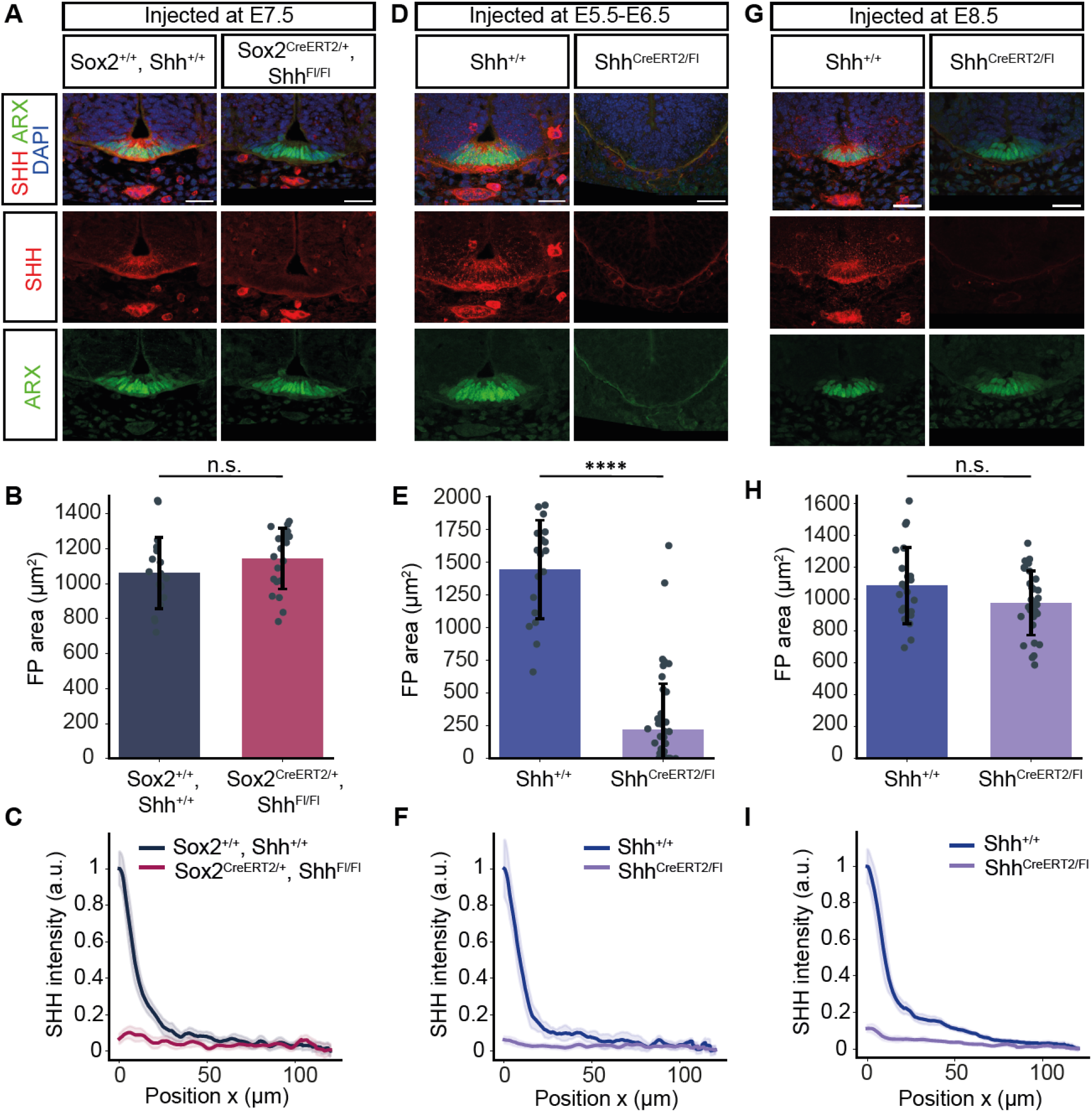
FP size is independent of FP-derived Shh, while the Shh gradient is maintained by the FP. **A**. Representative mouse brachial sections of E10.5 control and Sox2^CreERT2/+^, Shh^Flox/Flox^ embryos injected with tamoxifen at E7.5. Immunostaining as indicated. **B**. Quantification of the ARX-positive area in the experiment in A. Two-tailed *t*-test: not significant. **C**. Quantification of the SHH gradient in the receiving tissue in A. Mean profiles normalized to the maximum fluorescence intensity in the control condition with 95% CI (shaded) are shown. Number of sections: n=19 (control), n=21 (mutant). **D**. Representative images of E10.5 control and ShhCreERT2^CreERT2/Flox^ embryos injected with tamoxifen at E5.5 and E6.5. Immunostaining as indicated. **E**. Quantifications of the ARX-positive area in the experiment in D. Two-tailed *t*-test: *P* < 0.0001. **F**. Quantification of the SHH gradient in D, shown as in C. Number of sections: n=15 (control), n=48 (mutant). **G**. Representative images of E10.5 control and ShhCreERT2^CreERT2/Flox^ embryos injected with tamoxifen at E8.5. **H**. Quantifications of the ARX-positive area in the experiment in G. Two-tailed *t*-test: not significant. **I**. Quantification of the SHH gradient in G, shown as in C. Number of sections: n=22 (control), n=30 (mutant).

The insensitive class of networks is characterised by properties that can be experimentally tested. A key prediction of the model is that Shh input is required for FP production only at early developmental times, but is dispensable later on. To test these predictions, we used the Shh^CreERT2^ mouse line, in which a tamoxifen-inducible Cre is knocked into the Shh locus (Methods). We found that in Shh^CreERT2/+^ heterozygous embryos, the Shh gradient amplitude was reduced, but the floor plate appeared normal (Fig. S6A-C). We further crossed Shh^CreERT2/+^ mice to Shh^Flox^ mice to generate Shh^CreERT2/Flox^ embryos in which all Shh production from both the FP and notochord is deleted upon tamoxifen injection. We found that early deletion of Shh by injection at E5.5 and E6.5 resulted in a severe reduction of 85% of FP size compared to control wildtype littermates (Fig. 7D-F). By contrast, in embryos with late deletion by injection at E8.5, the FP size was not altered compared to controls (Fig. 7G-I). These results demonstrate that Shh input is dispensable for FP formation after E8.5 and confirms the model prediction.

A second prediction of the model is that Shh production is continuously required to increase the Shh gradient amplitude over time. To test this, we compared the Shh gradient shape in the Shh^CreERT2/Flox^ conditional mutants, in which all Shh production was eliminated at early or late time points. As expected, deletion of Shh at both time points resulted in a severe reduction (by 94% and 89%, respectively) in the Shh gradient amplitude (Fig. 7D, F, G, I). Consistent with the model, this indicates that Shh is turned over on a time scale of a day or smaller than a day. Crucially, this result confirms that new production of Shh is continuously required to maintain the gradient amplitude.

The model results further indicated that continuous flux of Shh from the notochord can contribute to the Shh gradient amplitude only if the magnitude of flux is high relative to the amount of Shh produced by the FP. Our experimental data indicate that the effect of Shh deletion from both the notochord and FP with Shh^CreERT2^ (Fig. 7I) compared to the deletion from FP alone with Sox2^CreERT2^ (Fig. 7C) have a similar effect on the Shh gradient amplitude. This indicates that in the embryo, after the initial rapid FP establishment phase, Shh flux from the notochord does not have a major contribution to the Shh gradient profile. Instead, our results suggest that once the floor plate is formed, it becomes the main source of the Shh gradient observed in the neural tube.

Overall, the experimental results support a mechanism of FP formation that is independent of FP-derived Shh. Our results demonstrate that FP formation requires only initial Shh input from the notochord and is not sensitive to loss of Shh signalling at later stages. By contrast, FP-derived Shh is essential for the increasing dynamics of the Shh gradient amplitude.

## Discussion

Despite its central importance to morphogen gradient formation, in many systems the dynamics of specification and growth of the morphogen source are poorly understood. Morphogen sources are not static, but change dynamically as organs grow and develop. In some systems, such as the Fgf signalling gradient in the presomitic mesoderm, morphogen production is distributed within the target territory and influenced by the overall growth and morphogenesis of the target (*37*). In many other systems, morphogens are produced in spatially restricted sources. This situation applies for instance to the Dpp gradient formation in the Drosophila wing disc, Nodal gradient formation in the zebrafish blastoderm, as well as roof plate and floor plate formation in the vertebrate neural tube. To form such sources, cells are specified to adopt source cell identity, and when this process is progressive, it leads to a shifting boundary between the source and target domains (*38*). In addition, ongoing tissue growth can influence the size of the source and thereby the overall morphogen production. How specification and growth contribute to the dynamic formation of producing domains and in turn to morphogen gradient shape is poorly understood.

Here, combining a dynamical model of Shh gradient formation in the vertebrate neural tube with experimental data, we dissected the contribution of tissue growth and cell identity specification to the formation the floor plate, which is a source of Shh production. Our model and experimental results showed that cell fate specification and growth contribute to the formation of the floor plate on different time scales. Initially, rapid cell fate specification regulated by the gene interaction network and Shh derived from the notochord establishes the relative floor plate size. Following this initial phase, the floor plate is passively expanded by tissue growth, resulting in scaling. Our findings are reminiscent of the specification of a Shh source, termed zone of polarizing activity, in the developing vertebrate limbs, which also occurs early in the development of the limb primordium and is followed by an expansion phase (reviewed in (*39, 40*)). The source of Hedgehog in the Drosophila wing disc is an extreme example of such a mechanism – it is specified from the onset of wing disc development and restricted by lineage to the posterior compartment (*41, 42*). These examples suggest that specification followed by scaling may be a common strategy used in the formation of discrete sources in developing tissues.

Consistent with this strategy, a key feature of the system that we identified is that the FP does not extend its own domain by positive feedback in which Shh production in induced in neighboring cells. This is in contrast to other systems, in which positive feedback expands the production domain. For instance the roof plate, the domain of BMP ligand production, transiently responds to BMP signaling and expands by positive feedback through the transcription factor Lmx1a (*43, 44*). Positive feedback is also involved in the formation of the Nodal morphogen gradient in zebrafish embryos and in mammalian gastruloids, in which the Nodal production domain expands to neighboring cells via a relay mechanism (*45, 46*). A relay mechanism has also been proposed for the Wnt gradient in planaria (*47*). While in the latter case the Wnt gradient relay has the potential to effectively convert the whole tissue into a producing domain, the Nodal source reaches a restricted size. Nodal signaling activity is limited via negative feedback that relies on the activation of the inhibitor Lefty as well as the co-receptor OEP (*48*). In the neural tube, Shh induces the expression of non-diffusible repressors of floor plate formation, such as Nkx2.2 (*49*), that restrict the floor plate size. Our model provides an opportunity to understand what distinguishes mechanisms of morphogen source formation that rely on positive feedback from those that do not in a quantitative systematic manner. We showed that networks that rely on positive feedback from Shh^FP^ to form and maintain the floor plate, correlate with weaker repression by N. By contrast, networks that are insensitive to positive feedback from Shh^FP^ require only weak activation of FP identity by Shh from the notochord and strong repressive interactions, as well as a larger contribution from uniform basal activators of FP identity.

Our results highlight the importance of basal activators for FP formation and raise the question of their molecular identity. Prior studies have shown that FoxA2, an early determinant of FP identity, can be induced by Nato3 in a Shh-independent manner (*50*). Sox2 transcription factors have been shown to influence the patterning of the ventral neural progenitor domains (*51*) and it is possible that they also affect FP formation. Furthermore, although the mechanisms of FP induction differ between species and anterior vs posterior positions along the body axis, in some cases FP identity is induced by Nodal signaling (reviewed in (*10*)), and more recently has been shown to be induced by uniform RA signaling in neural organoids (*52, 53*). These findings support the notion that Shh-independent activators of FP identity contribute to the formation of the FP.

Repressive interactions within the transcriptional network defining FP identity are also essential for floor plate formation. For practical reasons our model incorporates a simplified view of this network, however it is known that several additional interactions are involved in FP specification. FP development begins with an early specification step in which FoxA2 and Nkx2.2 are induced by Shh signaling and initially co-expressed in the same cells (*5*). Subsequently, FoxA2 induces Arx expression and represses Nkx2.2, Nkx2.2 induces FoxA2 and represses Arx (*5, 19, 49, 54*). It is possible that these additional interactions change the temporal kinetics of FP formation compared to our simplified model. Nevertheless, the time scales set by the stability of transcription factors in neural tube patterning have been shown to be relatively short, on the order of a few hours (*16, 44, 55*), suggesting that qualitatively the two phase dynamics of FP formation that we describe is likely to hold also in the context of a more complex gene network.

The Shh^FP^-insensitive mechanism allows neural tube patterning to initially depend on the notochord, thus coordinating the development of these two adjacent organs. At the same time, this mechanism allows the neural tube to become autonomous and independent of any long-term fluctuations or changes of Shh in the notochord, but rather ensures that the Shh source remains coordinated with the growth of the neural tube itself. Our modeling results indicate that the insensitive mechanism leads to scaling of the Shh amplitude with floor plate size and with tissue growth (Fig. 6B). Amplitude scaling of morphogen gradients also occurs in other systems and is sufficient to provide a large degree of pattern scaling (*56*). Nevertheless, it is important to note that measurements of the proliferation rate of the floor plate have shown that it is subject to domain specific regulation and is slower than in the neural progenitor domains (*17*). This implies that in reality the scaling of the floor plate with tissue size is regulated and occurs in a more complex manner. Slower proliferation of the FP alone would lead to underscaling, however, at later developmental stages, the overall growth rate of neural progenitors decreases due to neuronal differentiation and cell cycle lengthening. Thus, the relative floor plate area appears nearly constant (*17*). Nevertheless, the independent regulation of FP proliferation may have important developmental and evolutionary implications, allowing the Shh gradient amplitude and neural progenitor pattern to be tuned according to the species’ requirements.

The results from our biophysical model suggest that a Shh^FP^-insensitive mechanism of floor plate formation arises within given parameter ranges of the underlying network, without requiring any additional molecular mechanisms to maintain the FP unresponsive to Shh. However, in vivo, additional mechanisms are known to ensure that the FP is unresponsive. In mouse development, the notochord is in direct contact to the neural tube until approximately the 30 somite stage, while subsequently this contact is lost (*14*). It has been suggested that this may lead to an inability of the Shh derived from the notochord to continue spreading to the neural tube ((*14, 15*), also see (*57*)). Furthermore, the transcription of Gli transcription factors is repressed within the FP, which leads to an attenuation of Shh signaling in the FP (*5, 58, 59*). The tight regulation of FP sensitivity to Shh signaling highlights the functional importance of the insensitive mechanism for neural tube development. Consistent with this, it has been shown that the loss of responsiveness to Shh is a prerequisite for the elaboration and maintenance of FP identity (*5*). Our study provides a quantitative basis to further investigate the emergent properties of this mechanism and its influence on growth and patterning of the spinal cord.

## Acknowledgements

We thank Thomas Minchington and James Briscoe for comments on the manuscript. MZ, RH and MM were supported by a grant from the Priority Research Area DigiWorld under the Strategic Programme Excellence Initiative at Jagiellonian University. MZ and MM were supported by the Polish National Agency for Academic Exchange, and MZ received support from National Science Center, Poland, 2021/42/E/NZ2/00188. Work in the AK lab is supported by ISTA, the European Research Council under Horizon Europe: grant 101044579, and Austrian Science Fund (FWF): Grant DOI 10.55776/F78.

## Supporting Information

**Fig. S1.**
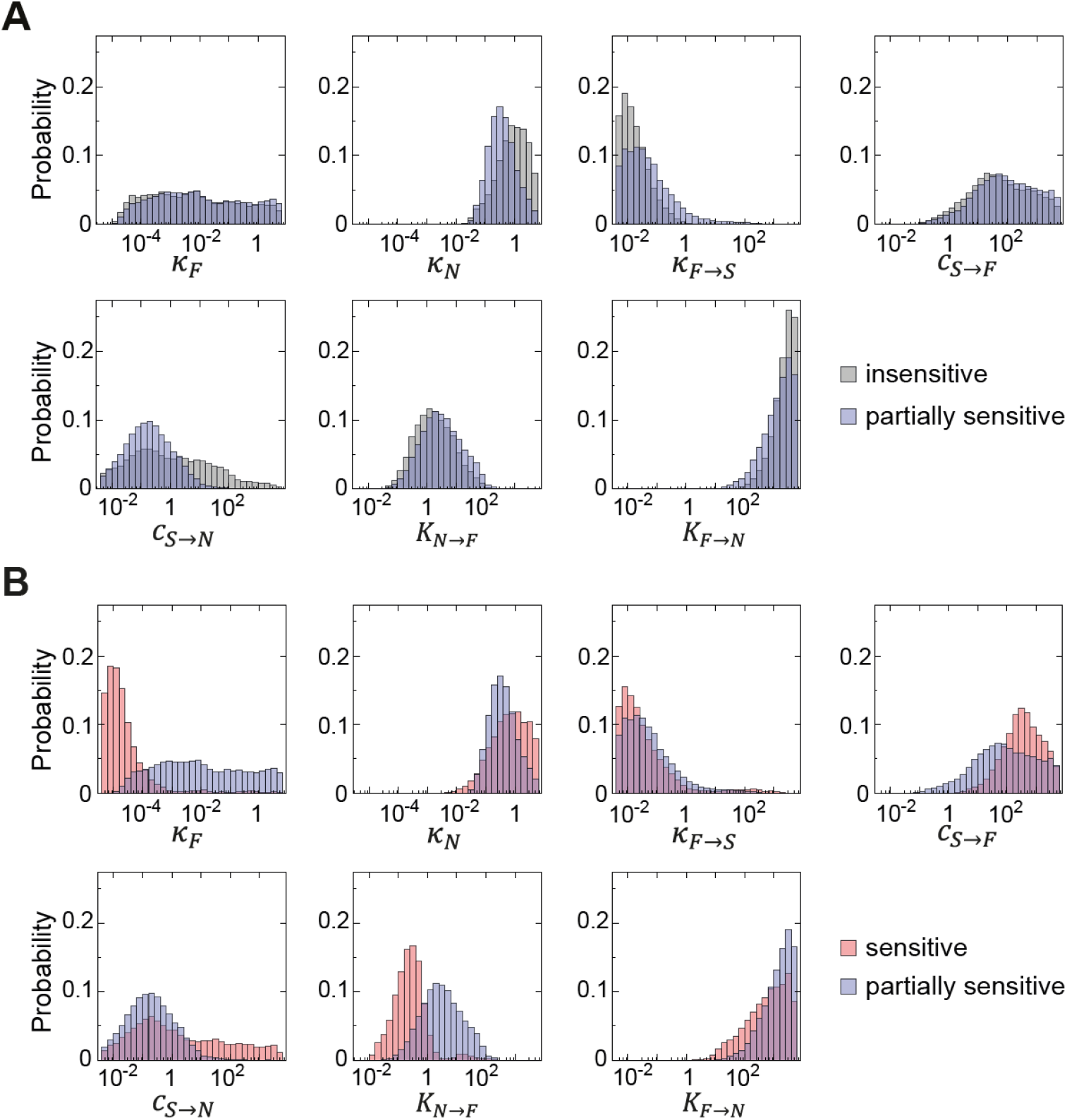
Model parameters for different mechanisms of FP formation. **A**. Distribution of model parameters for Shh^FP^-partially sensitive (blue) and insensitive (gray) solutions. In insensitive solutions the FP size does not change when Shh^FP^ is removed (*κ*_*F*→*S*_ = 0), whereas in partially sensitive solutions the FP size is decreased when Shh^FP^ is removed. **B**. The same as A, but for Shh^FP^-sensitive (red) and partially sensitive (blue) solutions. For sensitive solutions, the FP is completely lost when ShhFP is removed. A, B, insensitive (n = 66 827), partially sensitive (n=55 911), sensitive (n = 45 336).

**Fig. S2.**
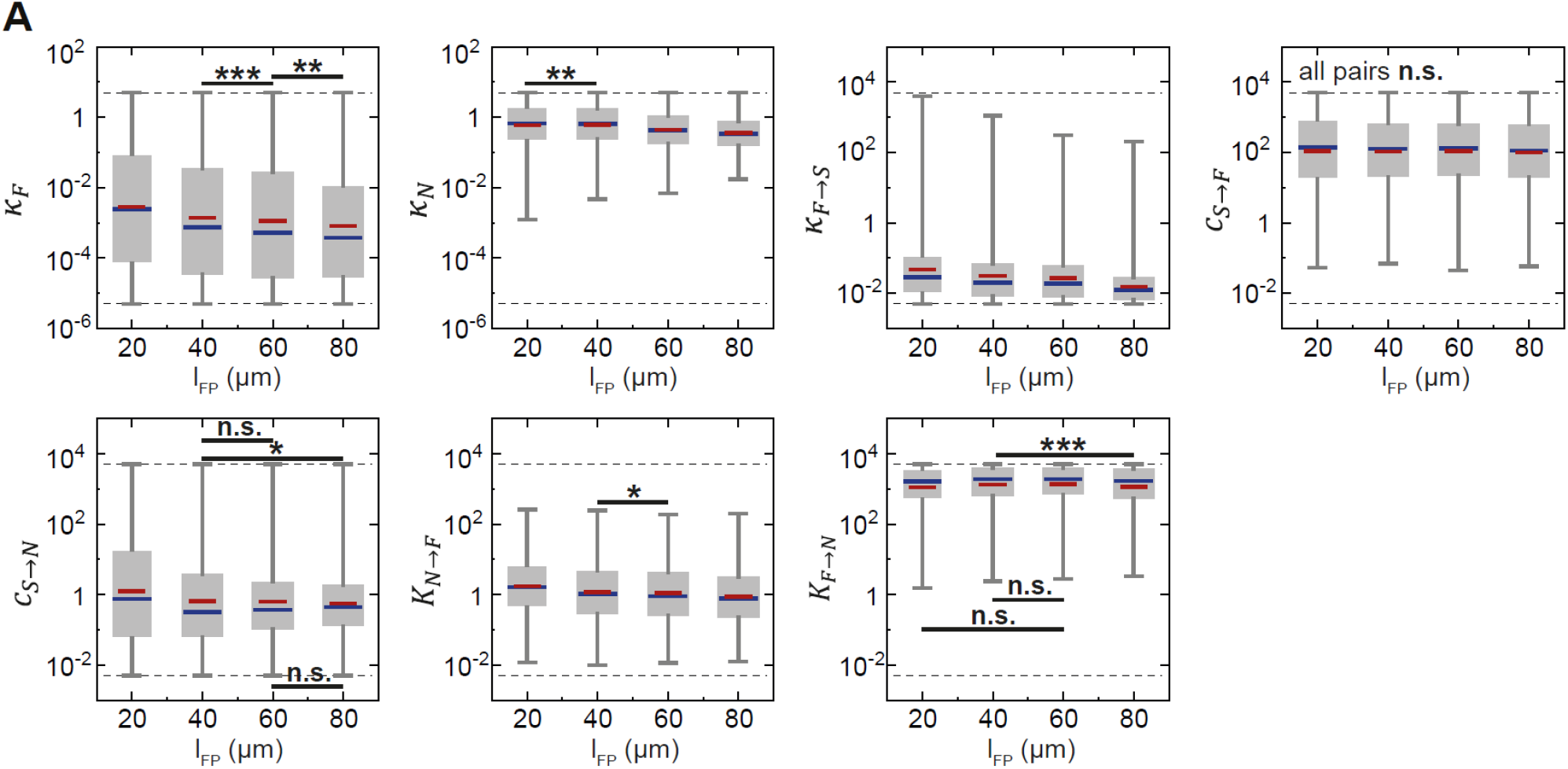
Dependence of FP size on model parameters. **A**. Distribution of model parameters for networks that yield the indicated final FP sizes at the end of the simulation (*L*_*end*_ = 400 μm at *t* = 60h). The boundaries of the allowed range of parameters in the screen are indicated with dashed lines. Box-whisker plots: 25–75 percentile (box), median (blue line), mean (red line), highest/lowest observation (fence). Pairwise comparisons two-tailed *t*-test: ns, not significant, *P* > 0.05; *, 0.05 ≥ *P* > 0.01; **, 0.01 ≥ *P* > 0.001; ***, 0.001 ≥ *P* > 0.0001; ****, 0.0001 ≥ *P*. All pairwise comparisons that are not indicated are ****. Sample sizes (number of solutions for given *l*_*Fp*_): 20μm (n = 23116), 40 μm (n = 12832), 60μm (n = 5254), 80μm (n = 1772).

**Fig. S3.**
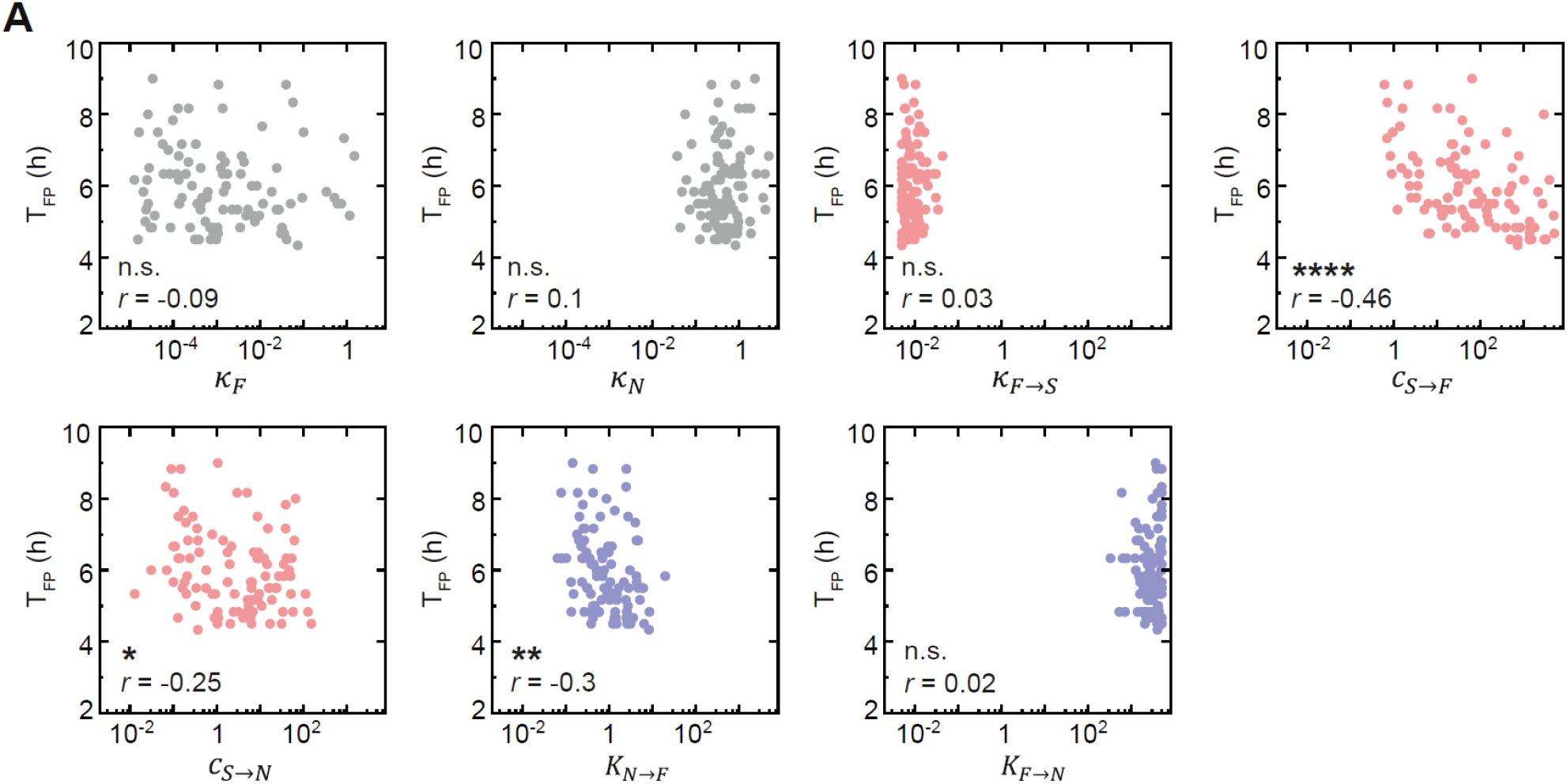
Dependence of FP formation time on model parameters. **A**. FP formation time for insensitive solutions for model parameters perturbed in the computational screen. The FP formation time is the time at which the FP reaches its final relative FP size. The Pearson’s correlation coefficient *r* between *T*_*FP*_ and log of parameter is reported. Pearson correlation test *P*-values: ns, not significant, *P* > 0.05; *, 0.05 ≥ *P* > 0.01; **, 0.01 ≥ *P* > 0.001; ***, 0.001 ≥ *P* > 0.0001; ****, 0.0001 ≥ *P*. n=100 solutions per parameter.

**Fig. S4.**
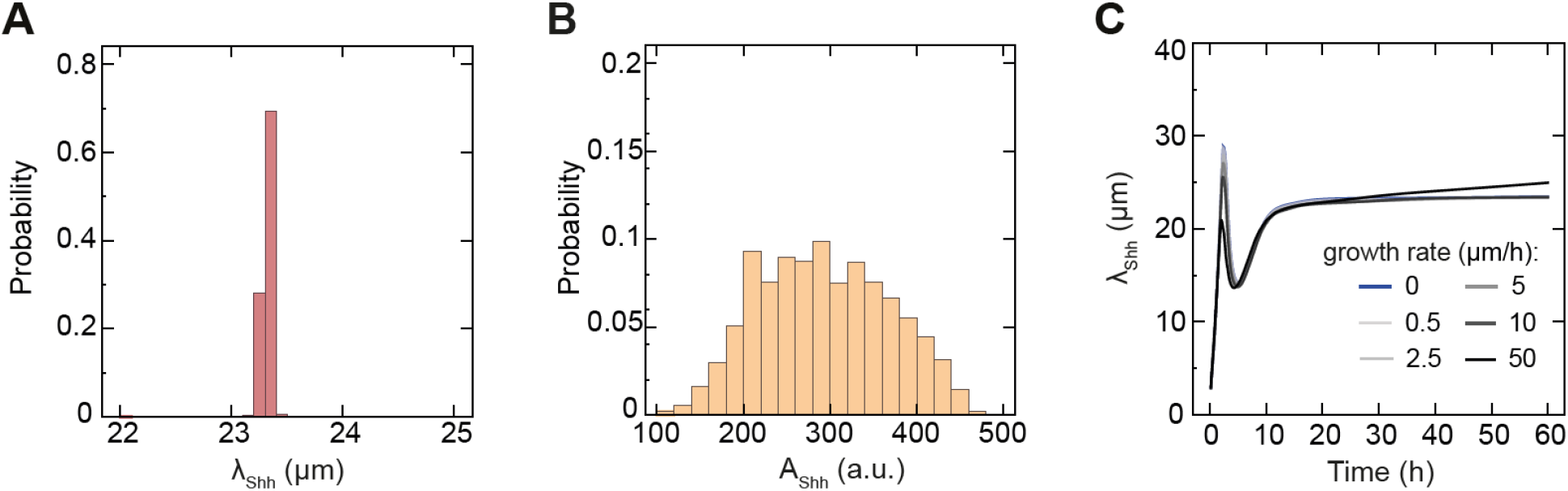
Shh decay length and amplitude in the computational screen. **A**. Probability distribution of Shh decay lengths in successful solutions of the computational screen. **B**. Probability distribution of Shh amplitudes. A, B, n = 169 979. **C**. Shh decay length for Shh^FP^-insensitive solutions as a function of time for different growth rates from *k*_*p*_= 0 μm/h to 50 μm/h (color coded as in the legend). The Shh decay length deviates from the value predicted by 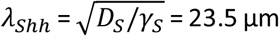 only for extremely fast growth. Sample size, n = 10 per condition, sampled every 10 min.

**Fig. S5.**
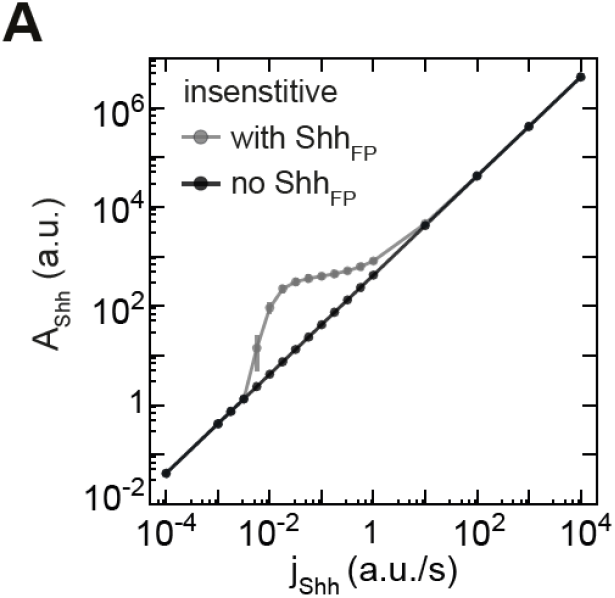
The Shh amplitude depends linearly on flux when FP-derived Shh is absent. **A**. Shh amplitude for large FP solutions with varied flux of Shh and no initial pulse of Shh. Without Shh^FP^ (black), the *A*_*Shh*_ increases linearly with *j*_*Shh*_. In solutions with Shh^FP^ present (grey), the amplitude is increased in the range of ∼0.01 ≤ *j*_*Shh*_ ≤ ∼1 a.u./s. The datapoints for the condition with Shh^FP^ are the same as in Fig. 6E. Sample size, n = 10 per condition, error bars SEM. For the condition with no Shh^FP^, the sampled points are identical for a given flux.

**Fig. S6.**
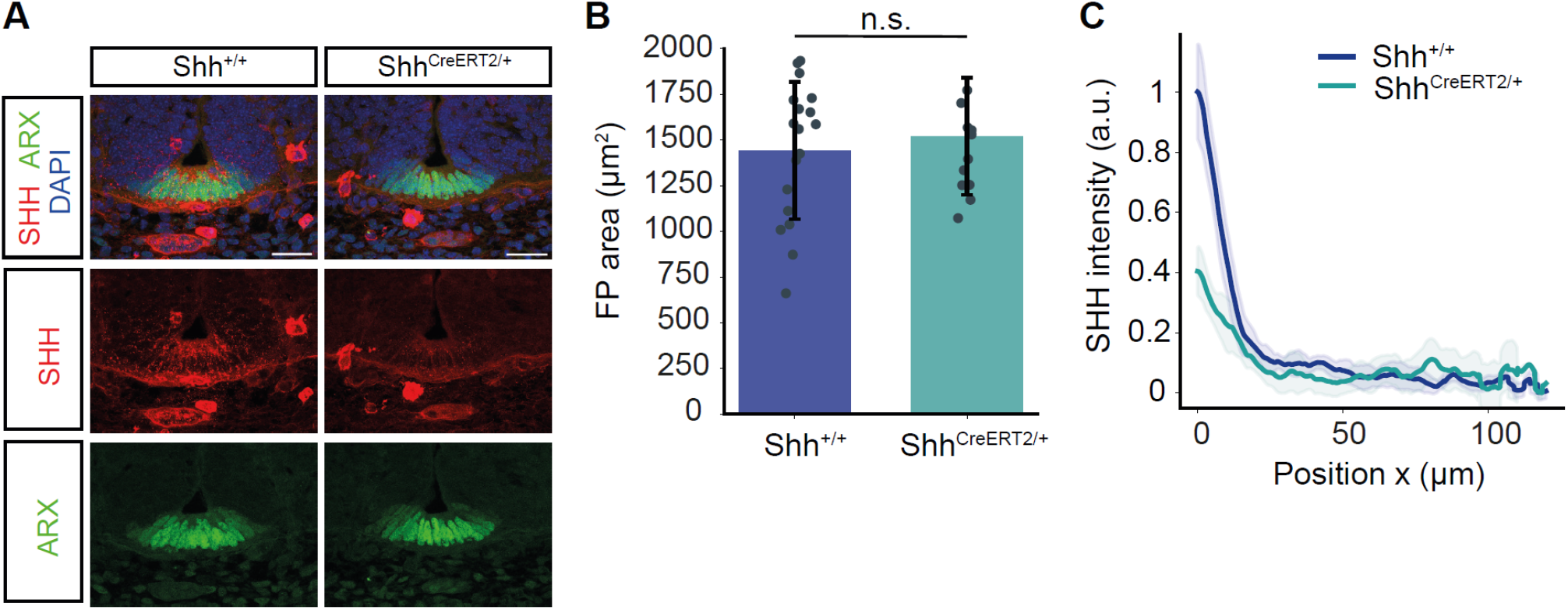
Loss of one copy of Shh reduces the Shh gradient amplitude, but does not affect FP size. **A**. Representative images of the ventral neural tube of E10.5 Shh^+/+^ and Shh^CreERT2/+^ embryos injected with tamoxifen at E5.5 and E6.5. The CreERT2 is knocked into the Shh locus, leading to a Shh null allele (see Methods for strain reference). **B**. Quantification of the ARX-positive area of the experiment in A. Two-tailed *t*-test: not significant. **C**. Quantification of the Shh fluorescence intensity in the receiving tissue of the experiment in A. Mean profiles normalized to the maximum fluorescence intensity in the Shh^+/+^ condition with 95% CI (shaded) are shown. n=15 sections for Shh^+/+^, n=13 sections for Shh^CreERT2/+^.

